# Single cell profiling reveals functional heterogeneity and serial killing in human peripheral and ex vivo-generated CD34+ progenitor derived Natural Killer cells

**DOI:** 10.1101/2022.01.24.477494

**Authors:** Nikita Subedi, Liesbeth Petronella Verhagen, Paul de Jonge, Laura Van Eyndhoven, Mark C. van Turnhout, Vera Koomen, Jean Baudry, Klaus Eyer, Harry Dolstra, Jurjen Tel

**Affiliations:** Laboratory of Immunoengineering, Dept. Biomedical Engineering, Eindhoven University of Technology, Eindhoven, Netherlands; Institute for Complex Molecular Systems, Eindhoven University of Technology, Eindhoven, Netherlands; Department of Laboratory Medicine - Laboratory of Hematology, Radboud Institute of Molecular Life Sciences, Radboud University Medical Center, Nijmegen, The Netherlands; Soft Tissue Engineering and Mechanobiology, Dept. Biomedical Engineering, Eindhoven University of Technology, Eindhoven, Netherlands; Laboratoire Colloïdes et Matériaux Divisés (LCMD), ESPCI Paris, PSL Research University, CNRS UMR8231 Chimie Biologie Innovation, Paris, France; Laboratory for Functional Immune Repertoire Analysis, Institute of Pharmaceutical Sciences, D-CHAB, ETH Zürich, Zurich, Switzerland

**Keywords:** Single cell study, Natural Killer cells, NK cell-based immunotherapy, Functional heterogeneity, Droplet based Microfluidics, Effector functions

## Abstract

Increasing evidence suggest that Natural killer (NK) cells are composed of distinct functional subsets. This multi-functional role displayed by NK cells have made them an attractive choice for anti-cancer immunotherapy. A functional NK cell repertoire is generated through cellular education, resulting in heterogeneous NK cell population with distinct capabilities to respond to different stimuli. The application of a high-throughput droplet-based microfluidic platform allows monitoring of NK cell-target cell interactions at single-cell level and in real-time. Through fluorescence-based screening of around 80,000 droplets, with different Effector:Target ratios, a fully automated image analysis allows for the assessment of individual killing events in each droplet over time. We observed a variable response of single NK cells towards different target cells and identified a distinct population of NK cells capable of inducing multiple target lysis, coined as serial killers. To meet the increasing clinical demand for NK cells several sources, such as umbilical cord blood (UCB), have successfully been explored. By assessing the cytotoxic dynamics, we showed that single UCB-derived CD34+ hematopoietic progenitor (HPC)-NK cells display superior anti-tumor cytotoxicity. Additionally, with an integrated analysis of cytotoxicity and cytokine secretion we showed that target cell interactions augmented cytotoxic as well as secretory behavior of NK cells. By providing an in-depth assessment over NK cell functions, this study provides crucial information on diversity and functional characteristics of peripheral blood NK cells and *ex vivo*-generated HPC-NK cells to develop and improve of NK cell-based cancer immunotherapy.

## Introduction

Natural killer (NK) cells are a subgroup of type 1 innate lymphoid cells capable of inducing cytolytic activity against virus-infected or cancer cells. Unlike other lymphocytes, these cells do not need prior antigen sensitization and induce rapid lysis of target cells upon identification without harming healthy tissue(Vanherberghen et al., 2013). Apart from cytotoxicity, NK cells also secrete immune-stimulating molecules that modulate the functions of other immune cells(Vivier et al., 2008). This functional versatility has enhanced the popularity to exploit NK cells in anti-cancer immunotherapy. More importantly, clinical effects have been demonstrated upon infusion in cancer-bearing patients(Hu et al., 2019; Liu et al., 2021). Recent developments in NK cell-based immunotherapy consist of different strategies, including the use of different cytokines to enhance autologous NK cells or to use allogeneic NK cells (both in their endogenous or genetically engineered form) as the mode of adoptive therapy. Even in allogeneic conditions, NK cell infusion is safe they match the criteria for the off-the-shelf immunotherapy(Fujisaki et al., 2009). A major hurdle is that only a small fraction of peripheral blood mononuclear cells comprises of NK cells, and thus generating them in sufficient numbers to meet clinical requirements is challenging(Fujisaki et al., 2009). Recently, other sources for NK cells, such as umbilical cord blood (UCB) and G-SCF mobilized blood, have therefore been successfully explored(Hoogstad-van Evert et al., 2017; Liu et al., 2020; Roeven et al., 2015) As a major advantage, UCB-derived CD34+ hematopoietic progenitor cells (HPCs) are more easily available and have several NK cell progenitors with fewer requirements for HLA matching compared to T cells.(Luevano et al., 2012). This makes UCB-derived HPCs a flexible and attractive source for NK cells(Hoogstad-van Evert et al., 2017). Upon expansion, these cells display significantly more cytotoxicity in bulk experiments compared to their blood-derived counterparts(Van der Meer et al., 2021).

In addition to their clinical relevance, it is also important to understand the functional potency of individual NK cells to optimize immunotherapeutic efficacy. Namely, NK cells exhibit a high degree of heterogeneity, both phenotypically and functionally(Ortaldo and Herberman, 1984; Smith et al., 2020; Yang et al., 2019). Mass cytometry-based data demonstrated how diverse the NK cell population can be by revealing over 100,000 different phenotypes within a healthy individual(Horowitz et al., 2013). Therefore, the overall function of NK cells results from combined efforts of multiple diverse individual cells. Although these cells lack genetic rearrangement and recombination, like T and B lymphocytes, this high variation shapes them into becoming a versatile cell type(Karo and Sun, 2015). Recent studies utilized single-cell RNA sequencing to profile NK cells transcriptomic and identified specific markers that could be clustered into different functional subpopulation(Crinier et al., 2021; Smith et al., 2020; Yang et al., 2019; Zhao et al., 2020). Furthermore, using a microwell-based single-cell platform Vanherberghen and coworkers identified rare serial killer NK cells with superior cytotoxic behavior which accounted for more than 50% of total lysis(Vanherberghen et al., 2013). Hence, single-cell-based tools lay the basis for a new era in dissecting heterogeneity within the NK cell compartment to address the functional ability of individual cells(Sinha et al., 2018).

Previously, we developed a high throughput droplet-based microfluidic platform to monitor the cytotoxic effector function of immune cells at single-cell level(Subedi et al., 2021). In this current study, we monitored around 80,000 droplets over 10 hours in real time to probe the phenotypical and functional heterogeneity of single human PB-NK cells and *ex vivo*-generated HPC-NK cells. Furthermore, we identified a cell population within these NK cell sources with superior serial killing ability. By pairing NK cells with distinct target cells, we demonstrate that NK cell-mediated cytotoxicity and cytokine secretion is a dynamic process restricted to a percentage of single NK cells equipped with this ability. By providing an in-depth assessment of NK cell functions, this research provides crucial information on the diversity and functional characteristics of PB-NK cells and HPC-NK cells to channel new avenues in NK cell-based cancer immunotherapy.

## Materials and Methods

### Cell isolation and culture

K562 cells were cultured in 1:1 (v/v) mixture of RPMI 1640 (Gibco, Catalog no. 22400089) and IMDM (Gibco, Catalog no.12440053) supplemented with 10% fetal bovine serum (FBS; Gibco) and 1% penicillin/streptomycin (PS; Gibco). THP-1 and Daudi cells were cultured in RPMI, supplemented with 10% FBS and 1% PS. All the cell lines were regularly tested for mycoplasma contamination. PB-NK cells were obtained from buffy coats of healthy donors (Sanquin) after written informed consent according to the Declaration of Helsinki and all experimental protocols concur to institutional guidelines. In short, peripheral blood mononuclear cells (PBMCs) were isolated from donor blood via density gradient centrifugation using Lymphoprep Density Gradient Medium (Stem cell). The NK cells were subsequently isolated using magnet-activated cell sorting (MACS) by negative selection using an NK cell isolation kit (Miltenyi Biotech, Catalog no. 130-092-657) following the manufacturer’s instructions. Cells were counted and purity was routinely assessed using flow cytometry by cell surface marker staining for 10 minutes at 4°C, using PE-CY7-labeled anti-CD56 (Biolegend, Catalog no. 362509), PE-labeled anti-CD16 (Biolegend, Catalog no. 302007), and PerCP-labeled anti CD3 (Biolegend, Catalog no. 300328) antibodies in 50 μL FACS buffer (2% FBS in PBS). The NK cells were identified as CD16^+^CD56^+^CD3^−^, and purity was on average 91%. Subsequently, isolated NK cells were encapsulated into droplets with K562 in presence of 1400 ng/mL IL2, as stimulant (Peprotech, Catalog no. 200-02).

### HPC-NK cell culture and expansion

Cryopreserved UCB CD34+ progenitor-derived HPC-NK cells from different donors were kindly provided by Dr. Harry Dolstra (Radboudumc, Nijmegen)(Van der Meer et al., 2021). The cells were thawed in medium with 71% Human Serum (HS; Sanquin), 0.03% DNAse and 0.1%MgCL_2_ and were washed (at 300 g for 15 minutes) after 10 minutes resting. The cells were then resuspended in NK MACS medium (Miltenyi, Catalog no. 130-107-209) supplemented with 10% HS and 1% NK MACS supplement (Miltenyi) at the concentration of 3 million cells/mL. 1.5 mL of the cell suspension was loaded into the 6 well plate and were supplemented by 50 ng/mL IL15 (Immunotools, Catalog no. 352310) and 0.2 ng/mL IL12 (Miltenyi, Catalog no. 130-096-704). On every second day, 1 mL NK MACS medium (with 10%HS, 1% NK MACS Supplement, 50 ng IL15 and 0.2 ng/mL IL12) was added to the cells, thus allowing them to expand for 7 days. All the assays were performed after the 6^th^ day of expansion. The culture was kept for 2 weeks after thawing.

### Microfluidic chip for droplet production

The three-inlet microfluidic device was developed following the protocols as described in Sinha et. al(Sinha et al., 2019). The microfluidic device was molded using an SU-8 photo resist structure on a silicon wafer and a commercially available polydimethylsiloxane silicone elastomer (Sylgard 184, Dow Corning), mixed with curing agent at the ratio 10:1 (w/w) and allowed to cure for 3 hours at 60^0^C. The surface of the Sylgard 184 was activated by exposure to plasma and sealed with a plasma-treated glass cover slide to yield closed micro channels. Channels were subsequently treated with a 5% (v/v) silane (1H,1H,2H,2H-Perfluorooctyltriethoxysilane; Fluorochem, Catalog no. S13150) solution in fluorinated oil (Novec HFE7500, 3M, Catalog no. 51243) and thermally bonded for 12 hours at 60^0^C. The dimensions of the microfluidic channels are 40 µm × 30 µm at the first inlet, 60 µm × 30 µm at the second inlet and the production nozzle, and 100 µm × 30 µm at the collection channel.

### Assembly of Droplet observation chamber

Glass microscopy slides (76 × 26 × 1 mm; Corning) were used as top and bottom covers (76 × 26 × 1 mm). Two access holes of 1.5 mm diameter were drilled in the top glass. Both slides were thoroughly cleaned using soap, water, and ethanol, and were exposed to air plasma (60 W) for 5 minutes. A cutout sheet of 60 μm thick double-sided tape (ORAFOL) was carefully placed above the bottom glass slide. Afterwards, the glass slides were stacked on top of each other, and the assembly was pressed using Atlas Manual 15T Hydraulic Press (Specac) for 5 minutes at 155°C at 400 kg per m^2^ pressure load. Next, two nano ports (Idex) were attached to the holes using UV curable glue (Loctite 3221 Henkel) which was cured under UV light for 5 min. Subsequently, the surface of the 2D chamber was treated with 5% (v/v) silane solution. Lastly, the chamber was dried, filled with fluorinated oil, and sealed until use. The chamber was reused multiple times and cleaned after each experiment by flushing fluorinated oil to remove droplets and was stored filled until the next use.

### Cell loading in microfluidic chip

Droplets were produced with a three-inlet microfluidic device. The protocol for loading cells in the microfluidic chips is described in Subedi et al(Subedi et al., 2021). The droplets of ∼50 µm diameter were generated using flow speeds of 30 µL/min for oil and 5 µL/min for each sample inlet. For serial killer experiments, droplets of ∼130 µm diameter were produced using flow speeds of 20 µL/min for oil and 5 µL/min for each sample inlet. The droplets were produced for around 5-10 minutes, thus generating 700,000 droplets in total. For the stability of droplets, 2.5% (v/v) Pico-Surf® surfactant (Sphere Fluidics, Catalog no. C024) was used in fluorinated oil.

### Bulk Activation Assay

NK cells were incubated at 1 million cells per 100 µL in PBA containing IFNγ Catch Reagent (Miltenyi, Catalog no. 130-090-443) and TNFα Catch Reagent (Miltenyi, Catalog no. 130-091-268) at 4°C for 20 minutes. Next, cells were washed and resuspended in RPMI cell culture medium supplemented with 2% HS, 1% PS, at 25,000 cells per 100 µL in U-bottom microwell plates together with stimulants (K562 cells at E:T 1:1 or IL2 50 ng/ml or K562+IL2 at above mentioned concentrations). The cells were incubated at 5% CO_2_ and 37 °C temperature for 4 hours.

### Single NK cell Activation Assay

NK cells were incubated at 1 million cells per 100 µL in PBA containing IFNγ Cytokine Catch Reagent and TNFα Cytokine Catch Reagent at 4°C for 20 minutes. Cells were then washed and resuspended in RPMI culture medium supplemented with 2% HS and 1%PS, at 2 million cells/mL for single-cell encapsulation. Next, the NK cells were encapsulated in 70 pL (∼50 µm) droplets together with stimulus (final concentration of K562 cells, 15 million cells/mL or IL2, 700 ng/ml or K562+IL2, earlier mentioned concentrations) loaded from another inlet. The concentrations of stimulus have been adjusted such that each single-cell received same absolute number of molecules as in bulk-based experiments. Droplet production and encapsulation rates were carefully monitored using a microscope (Nikon) at 10x magnification and a high-speed camera. The droplet emulsion was collected and covered with culture medium to protect droplets from evaporation. The encapsulated cells were incubated in Eppendorf tubes with a few punched holes to allow gas exchange, at 5% CO_2_ and 37 °C temperature. After 4 hours of incubation, the droplets were de-emulsified by adding 100 µL 20 v/v% 1H,1H,2H,2H-Perfluoro-1-octanol (PFO; Sigma Aldrich, 370533) in HFE-7500 and stained for FACS analysis.

### FACS-Antibody Staining

Cells were washed once with PBS and dead cells were stained with Zombie Green fixable viability dye (Biolegend, 423111), 1:10.000 in PBS, 50 µL) at 4°C for 20 minutes. Subsequently, cells were washed once with PBS and incubated with anti-human antibodies against CD3 (PerCP-Cy5.5, Biolegend), CD56 (BV510, Biolegend), CD16 (BV605, Biolegend), CD11b (PE-Cy5, Biolegend) CD27 (AF700, Biolegend) NKp46 (PE, Biolegend), NKg2A (PE-Dazzle 594, Biolegend), IFNγ (FITC, Miltenyi) and TNFα (APC, Miltenyi) at 4°C for 30 minutes. Next, the cells were washed twice with PBS buffer with 0.5% BSA and analyzed via BD FACS AriaII.

### Single NK cell cytotoxicity assay

NK cells and target cells were labelled with Calcein red (5 µM) (ATT Bioquest) and Cell tracker blue, CMAC (10 µM) (Invitrogen) respectively. The labelled NK cells and target cells were then loaded into droplet chip via different inlets at the concentration of 7 million cells/mL and 10 million cells/mL, respectively. The viability dyes Sytox Green (Invitrogen) and Cell Event Caspase-3/7 Green (Invitrogen) were loaded at the final concentration of 5 µM and 7 µM respectively along with the cells, and droplets were collected in the observation chamber. Droplets were generated at room temperature while collected into the observation chamber over a warm water bath. The immobilized droplets were incubated in a stage top incubator set at 5% CO_2_ and 37 °C. Image acquisition was performed at every hour interval for 10 h.

### Nanoparticle functionalization

50 µg of Paramagnetic nanoparticles (Bio-Adembeads Streptavidin Plus 300nm, Ademtech) were washed with 50 µg PBS(Gibco) using a magnet. The supernatant was removed, and the nanoparticles were resuspended in 990 µL PBS with biotinylated anti-IFNγ (Biolegend) antibodies and incubated for 30 min at room temperature while mixing. 10 µL Biotin was added with a final concentration of 1mM in the solution and incubated for 10 min at room temperature. The beads were washed again with PBS using magnets and resuspended in 5% Pluronic F-68 (Gibco) PBS solution and incubated for 30 min at room temperature. The beads were washed and resuspended in assay buffer containing RPMI 1640 (Gibco, life technologies), 5% Human Serum (HS) (Sanquin), and 25 mM HEPES (Gibco) and incubated for 10 min at room temperature. The nanoparticles were washed again and finally resuspended in the 100 µL of assay buffer containing fluorescently labeled AlexaFlora568-detection antibody for IFNγ (Biolegend).

When performing time-lapse experiments with cells, the final nanoparticle suspension contained 700 ng/ml IL-2 stimuli (Peprotech) and 10 μM Sytox Green (Invitrogen) as final concentration in drpplets. For experiments concerning the calibration curve and optimization steps, IFNγ (Peprotech) cytokine samples ranging from 0.001 - 100 nM were prepared in assay buffer. All calculations were made considering the final concentration inside the droplets.

### Image acquisition and analysis

Fluorescence imaging was performed using a Nikon Eclipse Ti2 microscope, using a 10X objective and mCherry, DAPI, and FITC/YFP filters every hour. The images were viewed using NIS Element and Image J. Automated Image analysis was performed using custom-made in-build MATLAB script (Mathworks), DMALAB (available with submission). The script generated a droplet mask that was overlaid onto the fluorescence images, and each droplet was analyzed separately. Over 80,000 droplets were analyzed using this script. The output received are in terms of droplet index, cell count, fluorescence intensity and dead cell count. Detailed description of image analysis script is provided in Subedi et al(Subedi et al., 2021). For the experiments with serial killers, microscopy images were analyzed manually.

### Statistics and software

The graphs were generated using GraphPad Prism 9.0.0. The results are expressed as mean ± SEM. Significant differences between two groups were analyzed by two-tailed unpaired Student’s t-test (Mann-Whitney test). P values < 0.05 were considered statistically significant. Flow cytometry data were analyzed using FlowJo X (Tree Star). FMO staining served as controls for gating strategy. For the gating strategy, the readers are referred to Supplementary Figure 1A.

## Results

### 1. Single-NK cell activation in droplets using K562 and IL2

To study the diversity within the NK cell compartment, we utilized droplets-based microfluidics to activate single NK cells with K562 cells and cytokines in an isolated environment with reduced paracrine interaction within cells. Similar to our previous work, NK cells were labelled with cytokine catch reagents for IFNγ and TNFα to allow for capturing and monitoring cytokine secretion by single-cells(Van Eyndhoven et al., 2021; Wimmers et al., 2018). After 4 hours of activation cells were retrieved from the droplets by breaking the emulsion with PFO and prepared the cells for downstream Flowcytometric analysis (Figure 1A). Similar to our previous studies with other cell types, the viability of NK cells after culturing in droplets was preserved (Supplementary figure 1B). We used pipette tips for loading cells in microfluidic chips to increase the probability of cellular encapsulation, and to achieve optimal cell pairing at a ratio of 1:1 in the oil-water droplets (∼70 pL)(Sinha et al., 2019). The cell loading concentrations were adjusted such that NK cells were optimally paired with K562 cells, and the percentage of activated cells reflected the cellular interaction. With optimal loading conditions we generated droplets containing 65% single NK cells co-encapsulated together with K562 cells (Figure 1B-D). In summary, our platform allowed us to probe NK cell activation at single-cell level and interaction with target cells.

**Figure 1.**
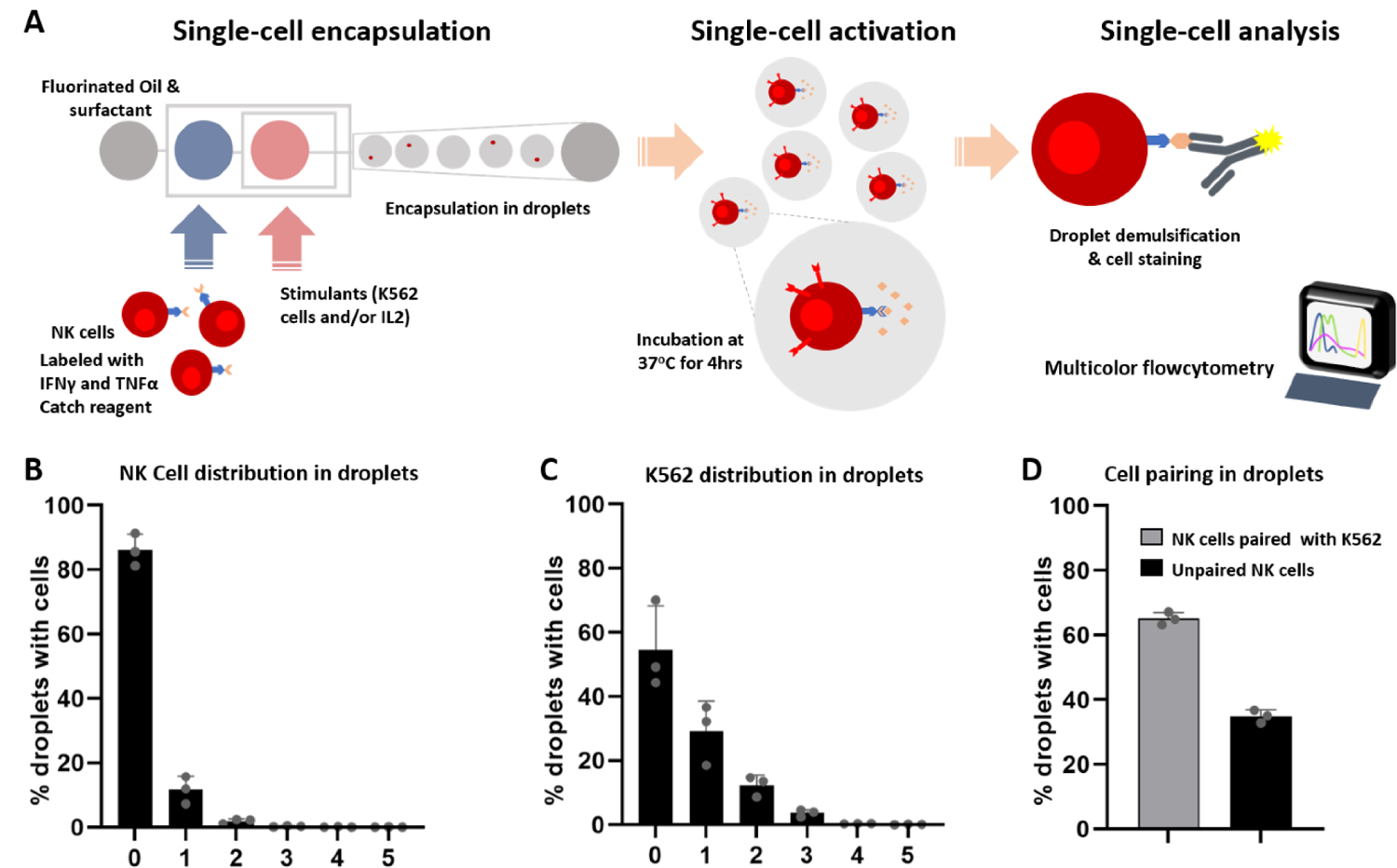
Experimental Schematics showing NK cell activation and cytokine secretion assay in droplet: **A.** Freshly isolated human peripheral blood NK cells were labelled with IFN γ and TNFα catch reagent and encapsulated together with stimulants (K562/IL2) into oil/water droplets using a 3-inlet flow-focusing droplet chip with 25 µm height. After 4 hours, the droplets were dissolved using PFO solution to retrieve the cells, which were thereafter labelled for FACS analysis. The distribution of cellular encapsulation of NK cells (**B.**) or K562 cells (**C.**) in the droplets. **D.** Cell pairing distribution in droplets. Results are shown as the mean ± SEM of 3 independent experiments with different donors.

### 2. Functional heterogeneity within peripheral blood NK cells - phenotypical correlation at different maturation stages

Studies associated stages of NK cell maturation with functional variations(Freud et al., 2017; Huntington et al., 2007). Here, we combined CD56 and CD16 with CD11b and CD27 to define different functional subsets within PB-NK cells. Even though CD11b and CD27 are generally not considered conventional markers for maturation in humans, their functional role has recently been documented(Fu et al., 2011; Vossen et al., 2008). Additionally, recent studies showed a correlation between murine maturation phenotypes and human phenotypes(Crinier et al., 2018). This prompted us to investigate these specific markers in association with human maturation stages. With our platform, we assessed the phases of maturation and characterized them into three distinct phenotypical subsets: 1. Tolerant NK cells (CD56^++^CD16^-^CD27^-^CD11b^-^), 2. Regulatory NK cells (CD56^++^CD16^-^CD27^+^CD11b^+^), and 3. Cytotoxic NK cells (CD56^+^CD16^++^CD27^-^CD11b^-^) (Figure 2A). We also identified an intermediate stage that showed dim expression of both CD56 and CD16 (CD56^+^CD16^+/-^ CD27^-^CD11b^+^), here described as pre-cytotoxic NK cells. The distribution of the different NK cell subsets was investigated in freshly isolated cells and analyzed after bulk and single-cell culture without addition of external stimuli. The composition within the population was dominated by mature NK cells, ∼80% CD56^+^ cytotoxic phenotypes (Figure 2B,C). Activating NK cell with IL2 induced a 2-fold increment in tolerant phenotypes (Figure 2D,F, Supplementary Figure 2) whereas interaction with K562 led to an increase in the cytotoxic subsets (Figure 2E,F, Supplementary Figure 2). For both single and bulk stimulated cells, we hardly observed the regulatory NK cell phenotype (Figure 2).

**Figure 2.**
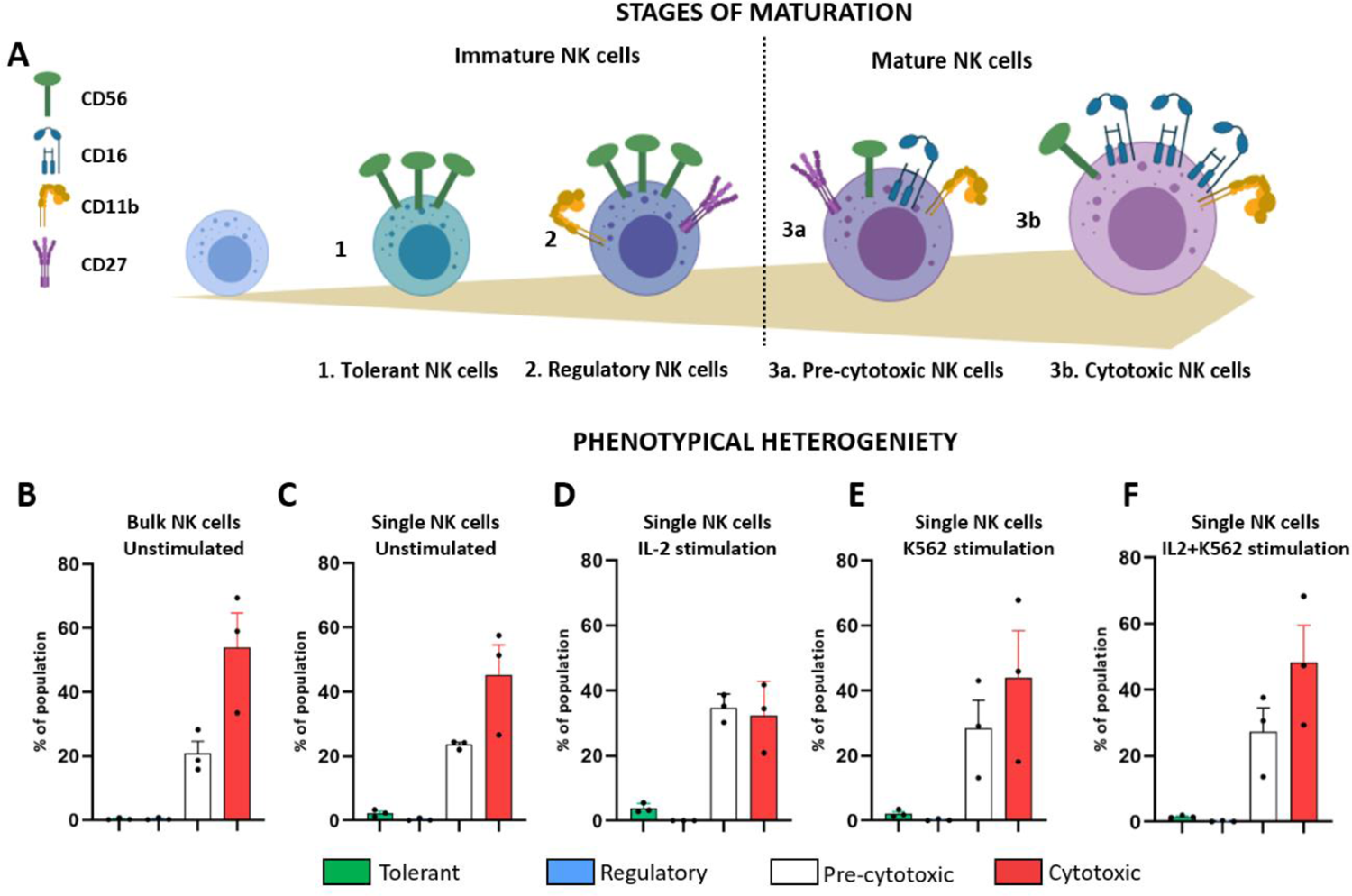
Phenotypical heterogeneity within peripheral blood NK cells and correlation at different maturation stages: **A.** Schematics of NK cell maturation versus different functional subgroups within the NK cell population. B-F, Graphs showing distribution of different NK cell population for different conditions upon incubation for 4 hours. **B.** Unstimulated NK cells in a traditional bulk-based assay. **C.** Unstimulated NK cells in droplets, **D.** Activation of single NK cells with IL2 in droplets. **E**. Activation of single NK cells with K562 in droplets. **F**. Activation of single NK cells with K562 and IL2. Green, blue, white, and red bars represent tolerant, regulatory, pre-cytotoxic and cytotoxic phenotypes respectively. Results are shown as the mean ± SEM of 3 independent experiments with different donors; different symbols (circle, triangle, and square) represent different donors.

Taken together, the overall composition of the PB-NK cell population with distinct subsets was not altered by external factors such as cytokines and cellular interactions. Even at single-cell level, the observed NK cell diversity was similar as was defined in earlier studies, thus confirming that droplets-based platforms are suited for studying NK cell heterogeneity.

### 3. Functional diversity of PB-NK cells at single-cell level

Next, we studied different functions of PB-NK cells at single-cell level to associate them with the respective phenotypes. IFNγ and TNFα secretion were included as functional markers for immune regulation, CD107a (degranulation marker) as measure for cytotoxicity and NKG2A and NKp46 as maturation markers(Abel et al., 2018; Schuster et al., 2016; Shigeru et al., 2008). The combined stimulation with IL2 and K562 induced the most secretion of IFNγ by all four defined subsets (Figure 3A). Regulatory and tolerant phenotypes (CD56^+++^) showed the highest percentage (approximately 10%) of cytokine secreting cells, however, the total number of events observed for these subtypes was very low (at most 2%). Therefore, compared to the absolute number of events, both of the cytotoxic phenotypes showed the largest number of positive cells upon combined stimulation. Both the regulatory and the cytotoxic phenotypes showed high TNFα secretion upon stimulation (Figure 3B). Notably, no difference in CD107a expression upon stimulation was observed with the cytotoxic phenotypes while the pre-cytotoxic phenotype dominated the CD107a expression thus showing an enhanced cytotoxic behavior (Figure 3C). Furthermore, pre-cytotoxic phenotype also formed a major component of the PB-NK cell compartment and expressed intermediate levels of NKp46 (Figure 3E). (generally expressed in mature NK cells). These findings strengthen the notion that this subset could be the immediate precursor of the cytotoxic population. As expected, the tolerant and regulatory phenotypes showed higher expression of NKG2A, while NKp46 was highly upregulated by more mature regulatory and cytotoxic phenotype (Figure 3D,E). Collectively, we showed functional variation among different PB-NK cell phenotypes upon stimulation at single-cell level indicative for heterogeneity within the NK cell compartment.

**Figure 3.**
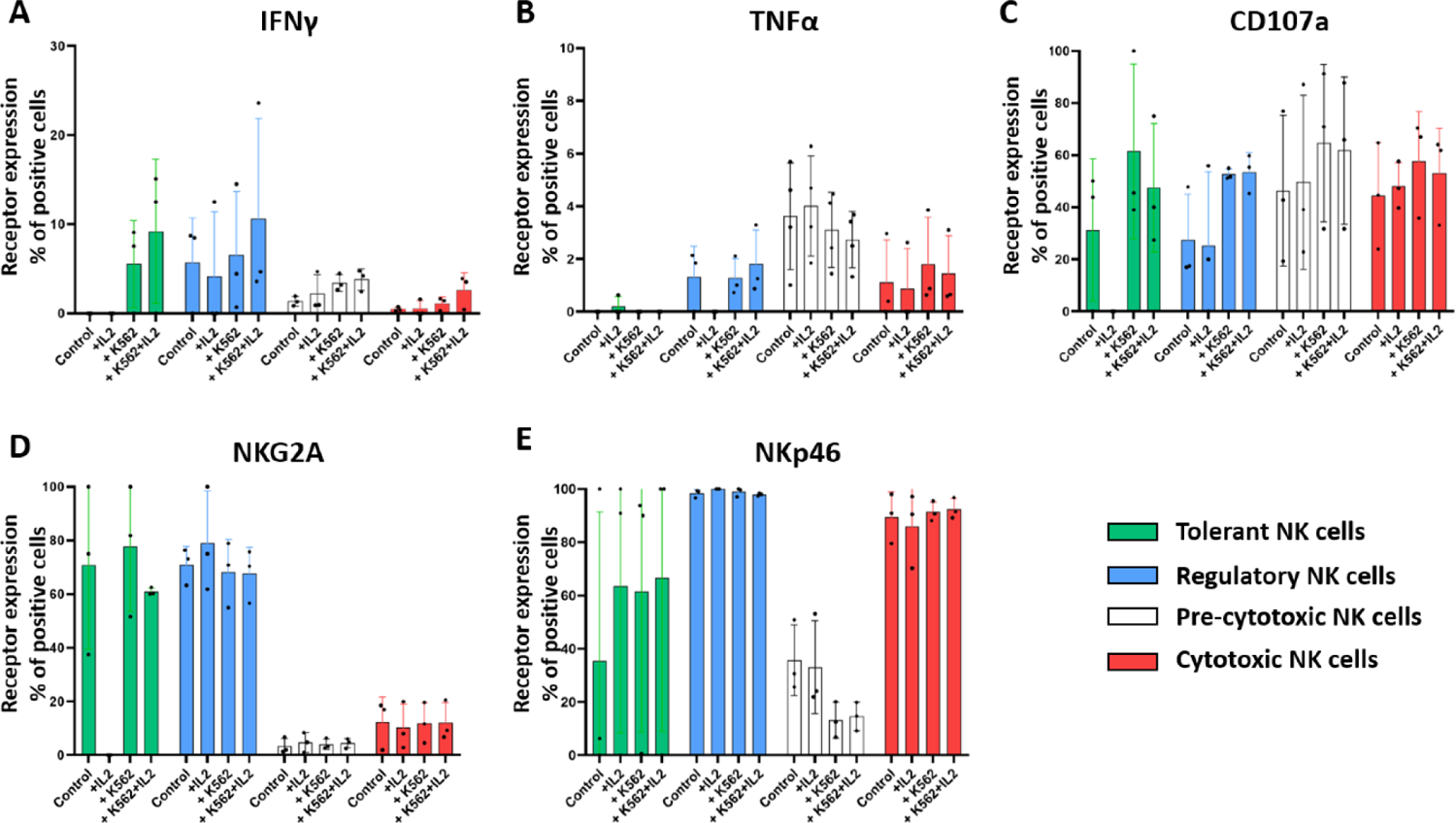
Functional diversity of NK cells at single-cell level: A-E, Graphs showing the percentage of NK cells population positive for **A.** IFN γ, **B.** TNF α, **C.** CD107a, **D.** NKG2A, **E.** NKp46 when stimulated in droplets with IL2, K562 or IL2 and K562 combination. Green, blue, white, and red bars represent tolerant, regulatory, pre-cytotoxic and cytotoxic phenotypes respectively. Results are shown as the mean ± SEM of 3 independent experiments with different donors; different symbols (circle, triangle, and square) represent different donors.

### 4. NK cell mediated cytotoxicity is a dynamic process and highly dependent upon target cell interaction

The interaction dynamics of single NK cells with different target cell types were closely monitored to improve the understanding of cytolytic function. Previously, we developed a droplet-based single-cell platform for high throughput and real time analysis of cellular cytotoxicity(Subedi et al., 2021). We used this platform to monitor over 80,000 droplets per experiment which approximately contained 3,500 droplets with a 1:1 NK cell: target cell ratio (Figure 4A). Individual droplets were monitored for 10 hours and subsequently analyzed with an automated script to identify possible cytotoxic events (Figure 4B). During the course of incubation, we observed around 3% target cell division (data not shown) however, all the E:T ratio were defined based on observation at t=0.

**Figure 4.**
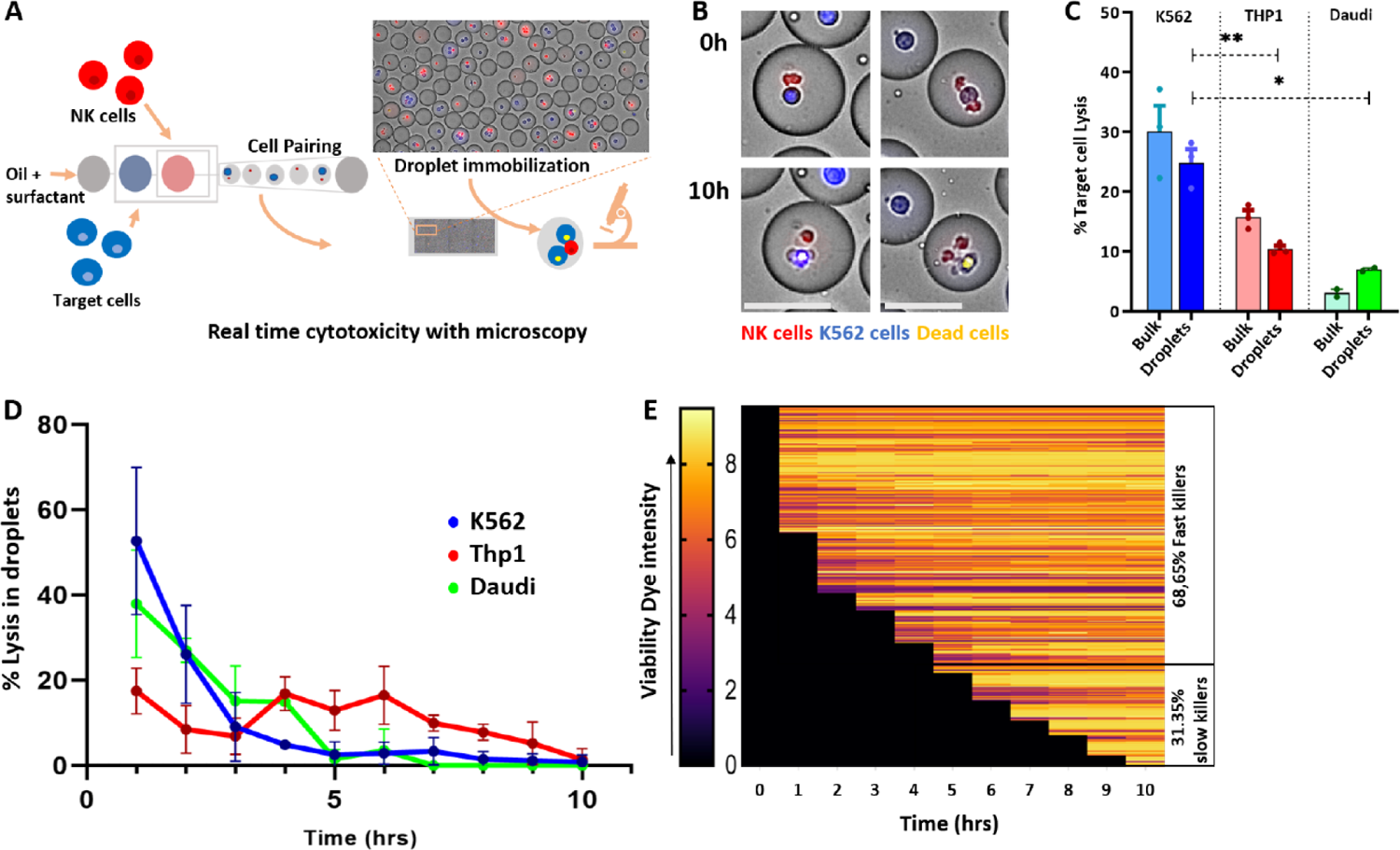
NK cell mediated cytotoxicity at single-cell level: **A.** Schematic representation of single-cell cytotoxicity. PB-NK cells were labelled with Calcein Red AM dye (red cells) and K562 cells with Cell Tracker Blue (blue cells) and paired together in 70 pL droplets in the presence of viability dye (Sytox green and Cell event Caspase 3/7; yellow cells). Cells were incubated at 37 °C and 5% CO_2_ for 10 h. **B.** Microscopic overview of NK cell-mediated cytotoxicity in droplets. Over the interval of 10 h, NK cells interacted with target cells inducing cytotoxicity. The dead cells were stained with the viability dye turning yellow. Scale bar = 50 µm. **C.** Bulk versus droplets cytotoxicity assay with K562 (Blue), THP-1 (Red) and Daudi cells (Green). Light and dark colors represent bulk and droplets experiments respectively. **D.** Graph representing the dynamics of cytotoxic events in droplets for different donors. The dynamics were determined for the killer fraction of NK cells, thus showing the percentage of the fast and slow killer population in NK cells. Blue, green, and red lines represent K562, THP1 and Daudi cells respectively; data analysis performed on E:T 1:1.; Each time point represents new events per hour. **E.** Heat Map showing the dynamics of cytotoxicity within an experiment. Each line represents individual cells. The graph does not include the droplets with 0, or 1 cell and droplets with dead cells at t = 0 h. Results are shown as the mean ± SEM of 3 independent experiments with different donors.

PB-NK cells showed a significant variation in the cytolytic function upon interaction with different target cells (Figure 4C). Where single PB-NK cells lysed around 25% (±2.25; n=3) K562 cells, only 10% (±4.318; n=3) of THP-1 and 8% (±0.271; n=3) of Daudi cells were killed by single PB-NK cells. We observed similar variation using a bulk-based cytotoxicity assay, thereby benchmarking our single-cell findings. For K562 and Daudi cells, most of the cytotoxic events were observed within the first four hours of cellular interaction, while lysis of THP-1 cells occurred at later time points (Figure 4D). This variation in lytic abilities of PB-NK cells could be linked with the differential expression of the MHC-I molecules expressed by different target cells (Supplementary figure 3).

Zooming in on the killing events revealed that 42% of NK cells (fast killers) induced killing as early as 1 hour. In total 69% NK cells were able to kill K562 cells during 4 hours of interaction (Figure 4E). The remaining 31% of cells (slow killers) only induced cytotoxicity at later time intervals. The observation of these early and late lytic events implies differential regulation in target cell recognition and the involvement of different cytotoxic mechanisms shown by different NK cells with regards to different target cells. Two distinct cytotoxic mechanisms were shown to act on different time scales, with rapid granule-mediated cell death and slower CD95-induced or TRAIL induced apoptosis(Prager et al., 2019). These striking differences in lytic ability observed between individual NK cells, together with their variation in killing behavior towards different target cells further strengthens the existence of heterogeneity within the PB-NK cell population.

### 5. Identification of rare serial killing events executed by NK cells using droplet-based single-cell platform

There exists a small fraction of NK cells endowed with an immense capacity to kill and handle most of the target cell lysis(Prager et al., 2019; Vanherberghen et al., 2013). These cells, also known as serial killers, can kill over 3 target cells consecutively and are a pursued phenotype for application in cancer immunotherapy. The fraction of serial killer NK cells is however relatively low and other than their superior killing ability, not much is known yet about them.

We therefore adapted the microfluidic platform by tuning the droplet size and cell loading concentrations and utilized the strength of our approach to study these potent serial killers in high throughput. With larger droplets (1.2 nL volume) we increased the fraction of droplets containing multiple target cells (≥3) with single NK cells to around 3% (242 droplets) (Figure 5A,B). Exploring different E:T ratios i.e., 1:1, 1:2, 2:1 and 3:1, variations in lysis of target cells was distinctly observed (Figure 5C). At 1:1 ratio, lower fraction of death in K562 cells was seen in bulk than compared to droplets at a 1:1 ratio. In contrast, at 1:2 ratio around 50% (±4.7; n=4) of paired K562 cells were killed by single encapsulated NK cells in the droplet and this was higher than observed in bulk-based measurements. When single K562 cells were paired with either 2 or 3 NK cells in droplets, the percentage of cell death increased and was somewhat higher than the corresponding bulk-based assay. By comparing with the significantly low killing events with unpaired K562 (in-droplet controls), we ruled out the possibility of spontaneous cell death in the droplets (Figure 5C).

**Figure 5.**
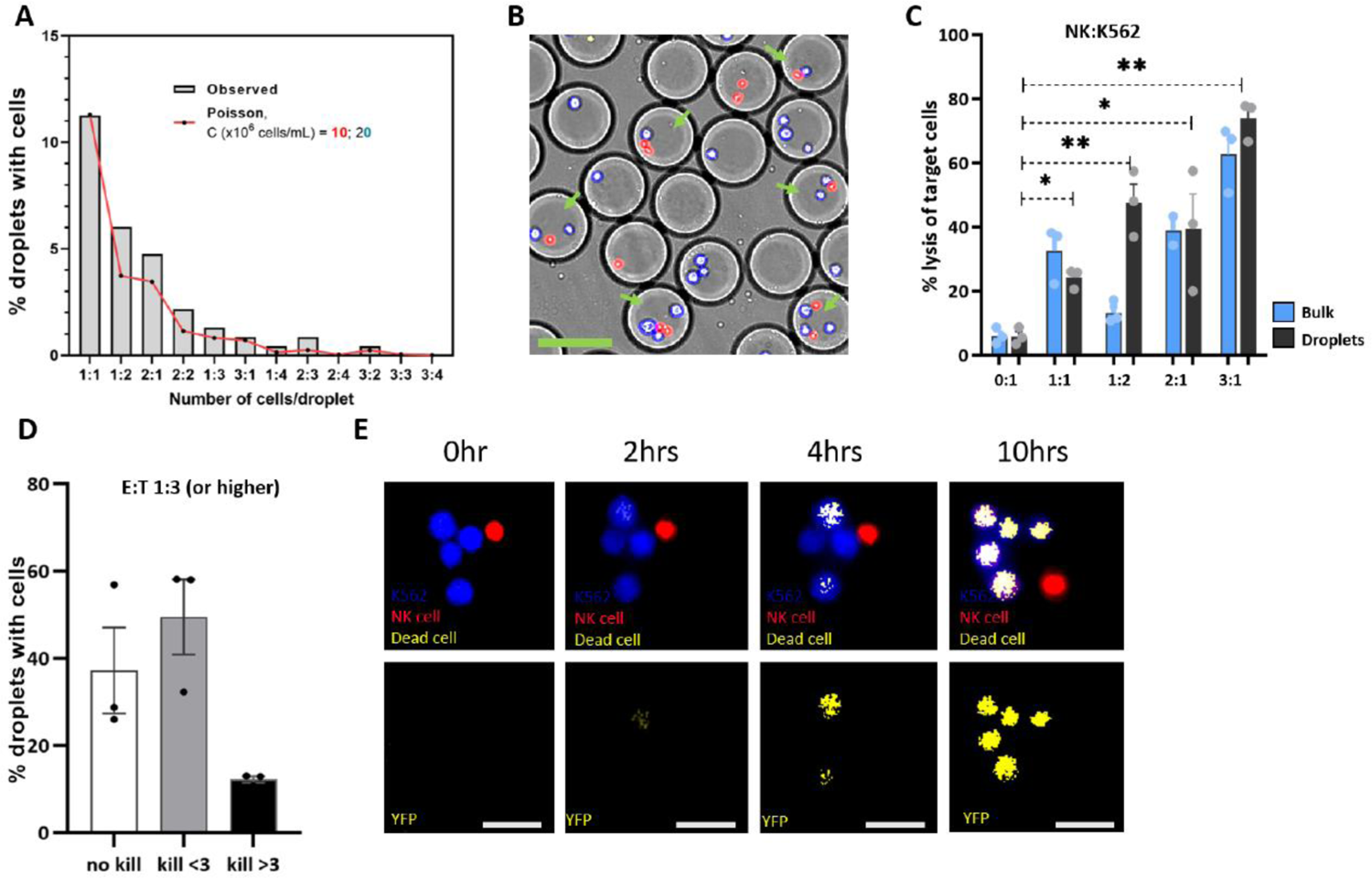
Adaptation of cytotoxicity platform to identify serial killer NK cells: **A.** Graph showing the experimental (grey bars) and predicted (red line) probability for different cell pairing ratios inside the 1.2 nL droplet. **B.** Microscopic overview of different E:T ratio for NK cells (red) and K562 (blue) observed inside the droplet. Scale bar=50 µm. **C.** Bulk (blue) versus droplets (black) cytotoxicity assay with K562 (at 0:1; 1:1; 1:2, 2:1, 3:1 E:T ratio, respectively. **D**. Graph depicting percentage of alive K562, <3 dead K562 and ≥3 dead K562 within the droplets containing more ≥3 or more K562 cells. **E.** Microscopic overview of a serial killing event in droplets over the period of 10 hours. red = NK cells; Blue= K562 cells; yellow= dead cells; scale bar= 50 µm

We defined serial killing activity if a single NK cell could consecutively lyse ≥3 target cells over the period of 10 hours. Zooming in on droplets with 1 NK cell and 3 or more target cells, we observed 12% (±0.736, n=3) serial killing events by PB-NK cells (Figure 5D,E, Supplementary figure 4). Additionally, we observed in 49% (±8.06, n=3) of droplets with less than 3 K562 cells target cell lysis and 37% (±9.68, n=3) droplets showed no target cell lysis. To ensure multiple cell encapsulation, we enhanced the droplet size which however resulted in later cellular interactions between NK and target cell, thus decreasing the percentage of positive cytotoxic events early upon incubation (Antona et al., 2020a).

Interestingly, using our droplet-based assay we segregated and created multiple conditions with high-throughput resolution to scrutinize the killing properties of NK cells. This enabled us to identify a rare serial killer subset in the PB-NK cell population where single NK cells are capable of lysing over three K562 cells. This is a first and important step to open the possibility of studying these highly searched for cell types.

### 6. Single *ex vivo*-generated HPC-NK cells display superior anti-tumor cytotoxicity

Given the potency and high translational efficacy of HPC-NK cells, we next sought to determine their tumoricidal activity. We observed an enhanced IFNγ and TNFα secretion by HPC-NK cells in droplets when paired together with target cells thus ensuring their activation in droplets Figure 6 A). In total 53% (±7.75; n=4) total lytic events was observed for these NK cells with an increasing trend of lytic events over the period of 10 hours (Figure 6B). HPC-NK cells were able to induce significantly higher lytic ability in comparison to PB-BK cells (Figure 6C). At different E:T ratios, both in bulk and at single-cell level, there was a significant increase in the percentage of cytotoxic events compared to unpaired target cells (Figure 6D). At 1:1, the cytotoxic event in both bulk (51%; ±7.4; n=4) and at single-cell level (54%; ±2.4; n=4) were similar, but with other E:T ratios the number of cytotoxic events increased in droplets despite the number of effector or target cells inside. A large fraction of NK cells with serial killing ability was identified in HPC-NK cells compared to PB-NK cells (Figure 6E and F). Of the total droplets with over 3 target cells, we observed around 34% (±3.2; n=4) HPC-NK cells had lysed over 3 target cells successively. We also observed a higher percentage of droplets with positive cytotoxic events (48%; ±4.0; n=4), while the droplets with no cytotoxic events remained at a minimum. In conclusion, we showed that *ex vivo*-generated HPC-NK cells are equipped with superior cytotoxic potential in comparison to PB-NK cells both at population as well as at single-cell level.

**Figure 6.**
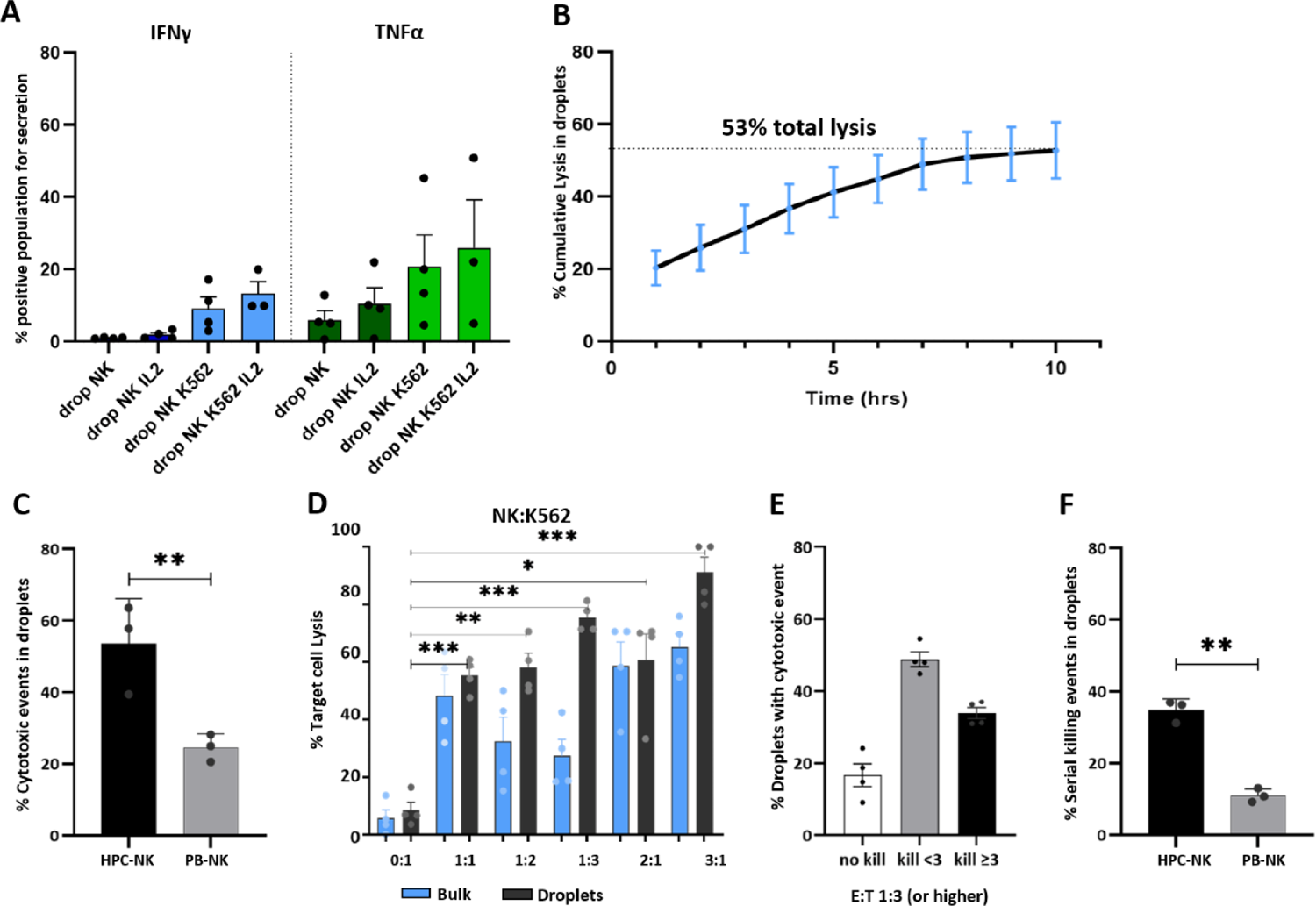
Functional assessment of HPC-NK cells in single-cell level: **A.** Activation of HPC-NK cells at single-cell level with different stimulants (IL2, K562 or IL2+K562) to measure the secretion of IFNγ (Blue bars) and TNFα (Green Bars). n=4 activation with IL2 and K562; n=3 activation with IL2 and K562**. B.** Graph representing the dynamics of cumulative cytotoxic events in droplets for different donor-derived HPC-NK cells. The dynamics were determined for the killer fraction of HPC-NK cells; data analysis performed all possible E:T. **C.** Comparison between cytotoxic events between UCB-derived (HPC) NK cells (Black bars) and PB-NK cells (grey bars); n=3. **D.** Bulk (blue) versus droplets (black) cytotoxicity assay with K562 (at 0:1; 1:1; 1:2, 1:3; 2:1, 3:1 E:T ratio, respectively. **E.** Graph depicting percentage of no dead K562, <3 dead K562 and ≥3 dead K562 within the droplets containing ≥3 K562 cells. F. Comparison between percentage of serial killers NK cells in HPC-NK cells (UCB-derived; black bars) and PB-NK cells (Grey bars); n=3. Results are shown as the mean ± SEM of 4 independent experiments with different donors (other than denoted differently).

### 7. NK and target cell co-encapsulation augments IFNγ secretion

To obtain insight in the distinct functional abilities of NK cells, we designed an in-droplet immunoassay to correlate two important functions: cytotoxicity and IFNγ secretion. To the best of our knowledge, we report for the first time on a droplet-based platform that allows monitoring both functions in single primary NK cells simultaneously in real time. A similar platform was developed by Antona et al. that also studied these functions in droplets, however their platform was designed for end-point based analysis and thus cannot address the temporal dynamics of cellular interactions(Antona et al., 2020b). We adapted the “In-drop sandwich immunoassay”, as described by Eyer.et al.(Bounab et al., 2020; Eyer et al., 2017) to allow the combinatorial investigation of IFNγ release and NK cell cytotoxicity in a time dependent manner. In essence, each droplet functions as a bio-nanoreactor containing NK cells co-encapsulated together with target cells, soluble viability dyes, functionalized magnetic nanoparticles, and detection antibodies in solution. Within a magnetic field, the nanoparticles form a uniform bead line. Thus, making each individual droplet a screening chamber in which cytotoxicity and secretion can be investigated together (Figure 7A,B). To validate our approach, we first generated two batches of droplets that both contained the magnetic capture beads and fluorescently labeled IFNγ detection antibodies and co-encapsulated either 0 nM IFNγ or 50 nM soluble IFNγ. During microscopy we clearly observed a strong positive signal for the droplet batch with soluble IFNγ (Figure 7C). Next, we integrated this technical advancement in our NK cell killing platform which yielded droplets with 7 different possible combinations (Figure 7D, Supplementary figure 5).

**Figure 7.**
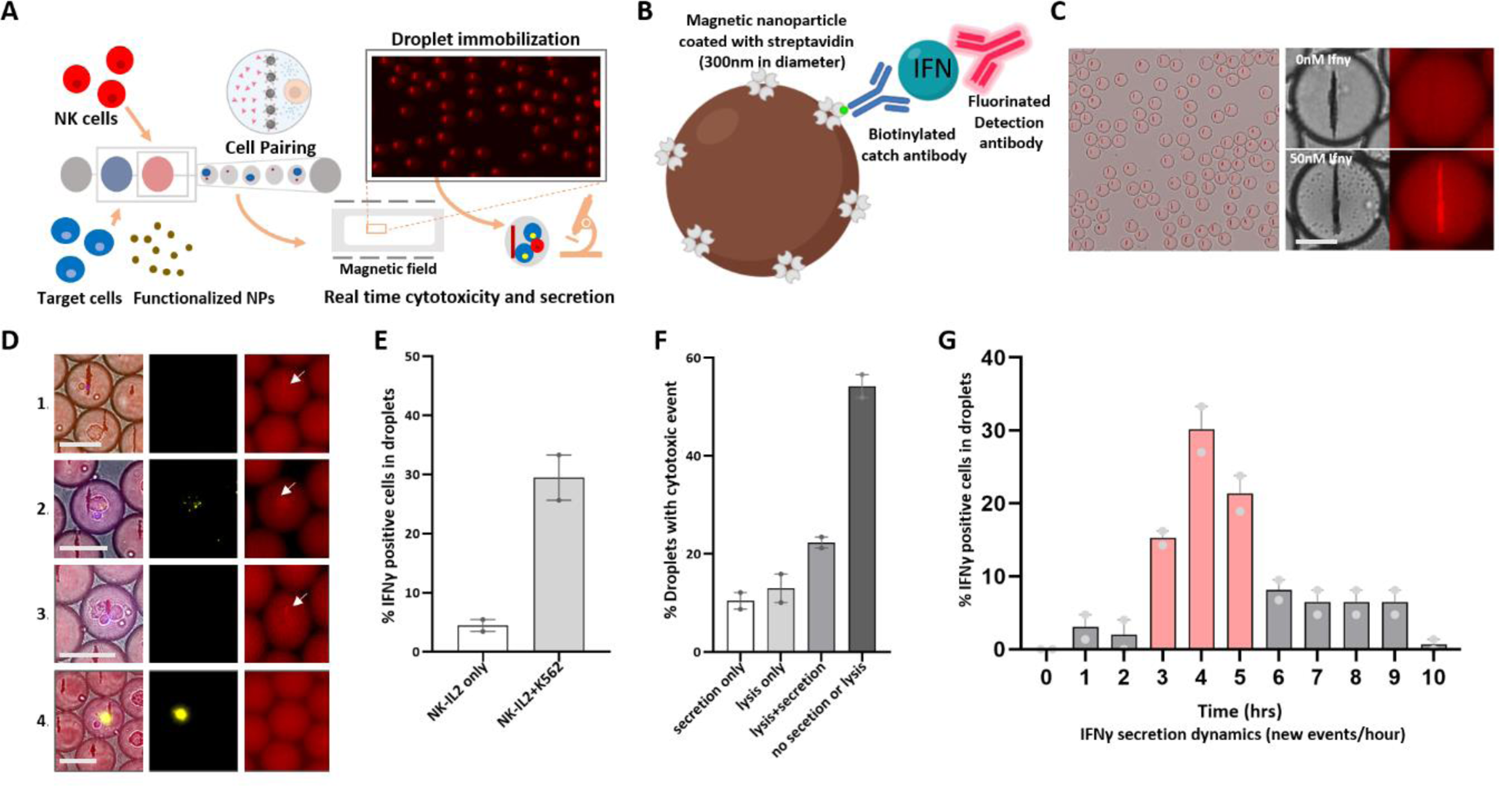
Combinatorial assessment of cytotoxic and secretory function of PB-NK cells upon target cell interaction: **A.** Schematics of in-droplet combinatorial assay. PB-NK cells were labelled with Cell tracker Blue (10 µM) and encapsulated together with target cells and nanoparticles mix at the concentration of 10 million cells/mL. Mixture of viability dyes (Sytox green and Cell event caspase3/7 green) was added together with the cells. Oil/water droplets were immobilized in an observation chamber in presence of magnetic field and monitored for 10 hours at 37°C. Each image was captured at an hour interval. **B.** The magnetic nanoparticle coated with streptavidin allowed binding of biotinylated catch antibody. When secreted IFNγ binds to the catch antibody, the freely floating detection antibody is relocated into the beads thus forming a fluorescent bead line. **C.** Microscopic overview of droplet with flourescent breadline (left) and zoomed in view of a droplet with 0 nM IFNγ (upper right) or with 50 nM IFNγ (lower right). Scale bar 50 µm. **D.** Microscopic overview of several conditions analyzed inside droplets; (1) Droplets with only NK cells that secreted IFNγ, (2) droplets with NK cells and K562 cells positive for both cytotoxicity and IFNγ secretion, (3) droplets with NK and K562 and positive only for IFNγ secretion, (4) droplets with K562 and NK cells positive only for cytotoxicity. Scale bar = 50 µm. **E.** Graph representing percentage of NK cells secreting IFNγ in droplets when incubated only with IL2 (white bars) or with IL2 and K562 (grey bars). **F.** Graph representing percentage of droplets with cell pair showing secretion only (white), lysis only (light grey); lysis and secretion (dark grey) and no secretion or lysis (black). **G.** Dynamics of IFNγ secretion by NK cells measured over the period of 10 hours. Each vertical bar represents independent events per hour. Results are shown as the mean ± SEM of 2 independent experiments with different donors.

To examine whether target cell-induced secretory and cytolytic function were associated with individual NK cells, we monitored the secretion dynamics of an NK cell upon pairing with target cells in real time. Approximately, 33% droplets were positive when paired with K562 while only 5% droplets without target cells showed positive IFNγ secretion (Figure 7E). Among the total number of droplets with E:T pairing, only 22% (±1.12; n=2) showed both cytotoxicity and secretion, 13% (±2.9; n=2) were positive only for cytolysis, and 10% (±1.6; n=2) for secretion only. To investigate how the dynamics of contact with target cells could regulate cytolysis and secretion in single NK cells, the droplets with positive events for both cytolysis and secretion were monitored closely. On average, an NK cell would already induce a lytic event within the first three hours of interaction (data not shown) while the secretion followed around 4.7 hours. The highest fraction of IFNy producing cells (∼67%) were observed positive within 3 to 6 hours (Figure 7G). Hence these results suggest that secretion follows the lytic event upon interaction with target cells, however the underlying mechanistic mode of action needs to be further explored.

## Discussion

Cellular heterogeneity within the NK cell compartment is well appreciated, however, how functional cellular properties are tied to this phenotypical diversity remains largely understudied. It is important to study at a single-cell level in a noise-free environment to exclude juxtacrine or paracrine interactions to fully comprehend NK cell diversification and their ability to induce different effector functions. Therefore, an optimal experimental approach requires both stimulation and analysis with single-cell resolution. By activating NK cells in picolitre size droplets, we ensured that external noise is reduced, and that observed cellular responses reflect intrinsic behavior.

Using our microfluidic platform, we present the functional assessment of *ex vivo*-generated HPC-NK cells and compared them with peripheral blood NK cells at single-cell level. We successfully demonstrated that HPC-NK cells, upon expansion with cytokines, are significantly more cytotoxic than primary NK cells isolated from peripheral blood. Furthermore, we showed that HPC-NK cells harbor a larger pool of serial killers which is consistent with the percentage of serial killers in HPC-NK cells identified in a recent study using a microwell-based platform(Van der Meer et al., 2021). Interestingly, for all E:T ratios tested (except for 1:1), we observed a higher percentage of cytotoxicity similar or higher than bulk-based analysis. These results suggest that the confinement within droplets enhances the probability for NK cells to interact with a target cell compared to a crowded microwell-based setup(Antona et al., 2020a). Similar cytotoxic events at different E:T ratio, for example similar percentage of lysis with 2:1 and 1:2 (or 3:1 or 1:3) was observed. This suggests that NK cells within a small group (as inside the droplets), operate independently to mediate the lysis of a single target cell and do not show cumulative cytolytic effect by cooperating with neighboring NK cell as in bulk based co-culture(Yamanaka et al., 2012a).

Serial killers have been studied previously for population-based IL-2-activated human PB-NK cells(Bhat and Watzl, 2007; Christakou et al., 2013; Vanherberghen et al., 2013). However, it is still not studied how NK cell activation at a single-cell level affects the phenomenon. We identified 12% serial killing events where a single NK cell (upon single-cell activation) could lyse ≥3 target cells consecutively. The percentage of serial killers identified in our study is higher than what had been observed in resting NK cells, as shown by Guldevall et.al., suggesting that IL2 can enhance the serial killer behavior in NK cells(Bhat and Watzl, 2007; Guldevall et al., 2016). In line with earlier studies, around 25% of NK cells showed positive cytotoxic events while remaining other NK cells did not induce target cell lysis(Guldevall et al., 2016; Subedi et al., 2021; Vanherberghen et al., 2013). In contrast to the work described by Sarkar et.al., we did not observe 100% NK cell-mediated killing in droplets. We believe 100% killing in earlier study could be due to the characteristics of the utilized dye being actively pumped out by the target cell(Sarkar et al., 2017).

The FACS based data from single-cell activation showed an augmented secretory activity of CD56^+^ phenotypes (both cytotoxic and pre-cytotoxic) upon interaction with their target cells(An et al., 2017). We further investigated the correlation between target interaction induced cytotoxic event and secretory behavior of individual NK cells by incorporating an innovative in-droplet sandwich immunoassay together with our in-droplet cytotoxicity assay.

This combinatorial platform provides a unique ability for high throughput monitoring of cytokine secretion in real time together with cytotoxicity at single-cell level. To this end we identified 74% of total lytic events in droplets that showed positive secretory function as well. In this way we demonstrated a correlation between cytotoxic and secretory function of NK cells, which others were not able to find(Antona et al., 2020b; Yamanaka et al., 2012b). We believe that this discrepancy is explained by a short experimental protocol of 4 hours leading to a missed cytolytic fraction that also test positive for IFNγ.

The dynamics of NK cell-mediated cytotoxicity are dependent on several factors such as the maturation state of NK cells, phenotypical variation based on the expression of several surface molecules, and lytic content of NK cells(Sarkar et al., 2017). Besides these NK cell-specific factors, the expression of NK-responsive factors in target cells also determines the nature and speed of cell death. In our study, we presented the comparison of NK cell-mediated cytotoxicity with two leukemia (K562 and THP-1) and one lymphoma (Daudi) cell line, known to show different sensitivity towards NK cells(Müllbacher and NJ; Zarcone et al., 1987). Along with the number of cytotoxic events, the differences were also observed in the timeline of the cytotoxicity. This variation could be linked to the surface arrangements of these cells (expression patterns of different activating and inhibitory molecules) which eventually leads to activation of different killing mechanisms. In agreement with the literature, we observed upregulated expression of HLA molecules by all cell types(Fayen and Tykocinski, 1999; Ramirez et al., 1992; Tsuchiya et al., 1980). NK cells lyse K562 target cells primarily by delivering perforin/granzyme-loaded cytolytic granules into the lytic synapse. However, the lysis of THP-1 cells had been found to be more dependent on cytokines, such as IFNγ that could lead to increase in ICAM-1 molecule upon exposure(Lehmann et al., 2001; Wang et al., 2012). NK cells also kill THP-1 cells by forming nanotubes that generally occurs after certain hours of interaction(Chauveau et al., 2010; Wang et al., 2012). Involvement of all these different pathways lead to later killing of THP-1 cells compared to K562.

Our research puts emphasis on unraveling the complex functional and phenotypical heterogeneity within the NK cell population. This research provided an integrated analysis of NK-target cell interactions and its implications on the cytolytic and secretory behavior of single NK cells. By adapting the droplet-based cytotoxicity platform, we identified rare serial killers, thus channeling exciting ways for easy identification and study of these rare cell types in future. Furthermore, with functional assessment of *ex vivo*-generated HPCNK cells, one of the important sources of adaptive NK cell therapy, we probed heterogeneity in these cell types. We believe that our data on functional heterogeneity underlying NK cell population, both peripheral NK cells as well as CD34+ HPC-derived NK cells, provides valuable contributions towards developing and elevating efficacy of NK cell-based cancer immunotherapy.

## Author Contributions

NS, LVV and, JT designed the study. NS, LVV and, VK performed the experiments. KE and NS developed the observation chambers. MVT and KE wrote the script. NS, LVV, VK analyzed the data. NS, LVV, LVE and, JT wrote the article. PJ, HD, KE and, JB verified the findings and provided resources for the project. JT supervised the research, verified the findings, validated the manuscript, and acquired the funding.

## Acknowledgments

These results are part of the project that has received funding from the European Research Council (ERC) under the European Union’s Horizon 2020 research and innovation programme (Grant agreement No.802791). Furthermore, we acknowledge generous support by the Eindhoven University of Technology.

---

**Supplementary Figure 1:**
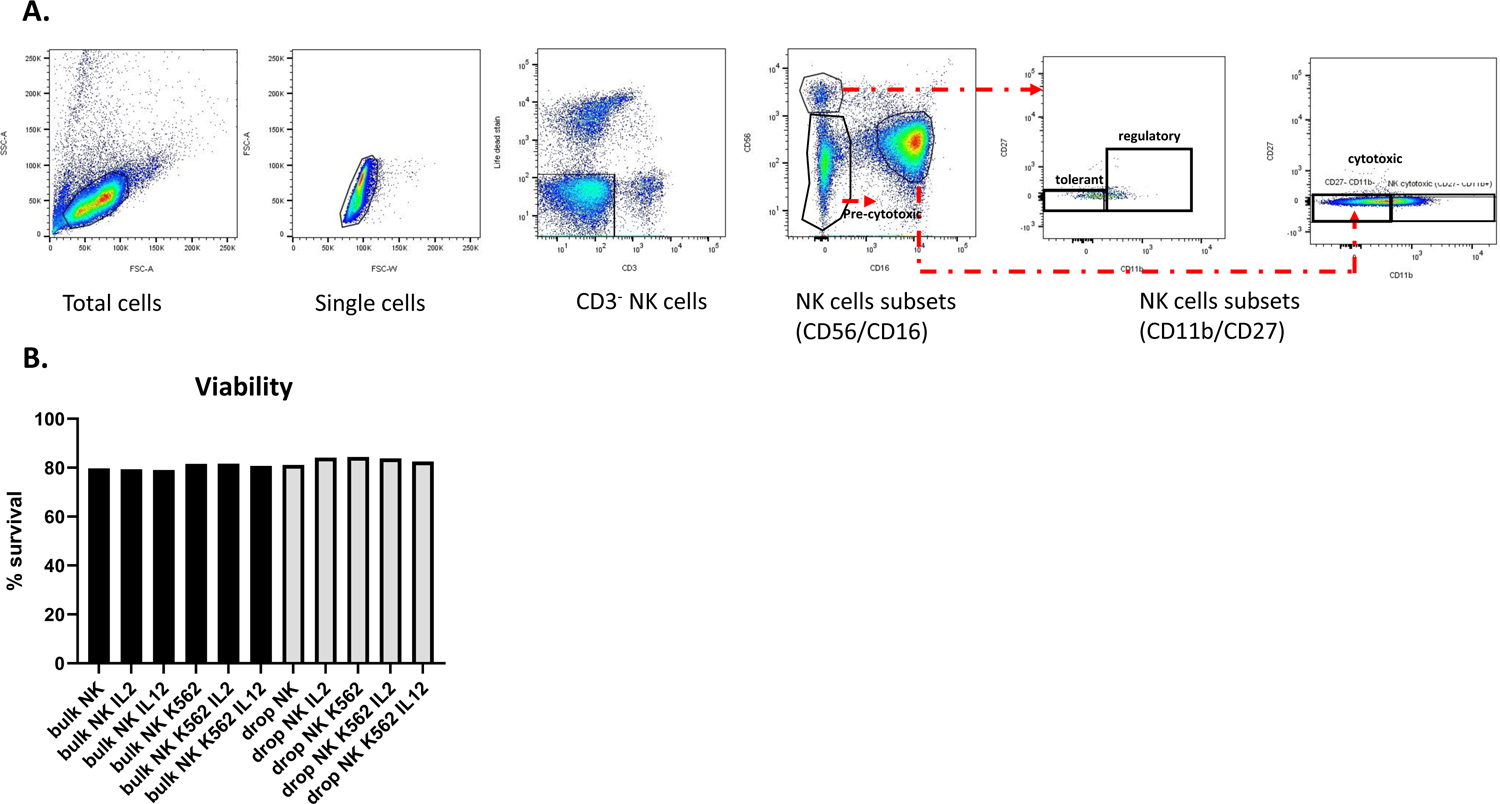
**A.** Gating strategy for NK cells to identify different subpopulation based on CD56/CD16 and CD11b/CD27. **B.** Graph showing the viability of NK cells in bulk and droplets after activation.

**Supplementary Figure 2:**
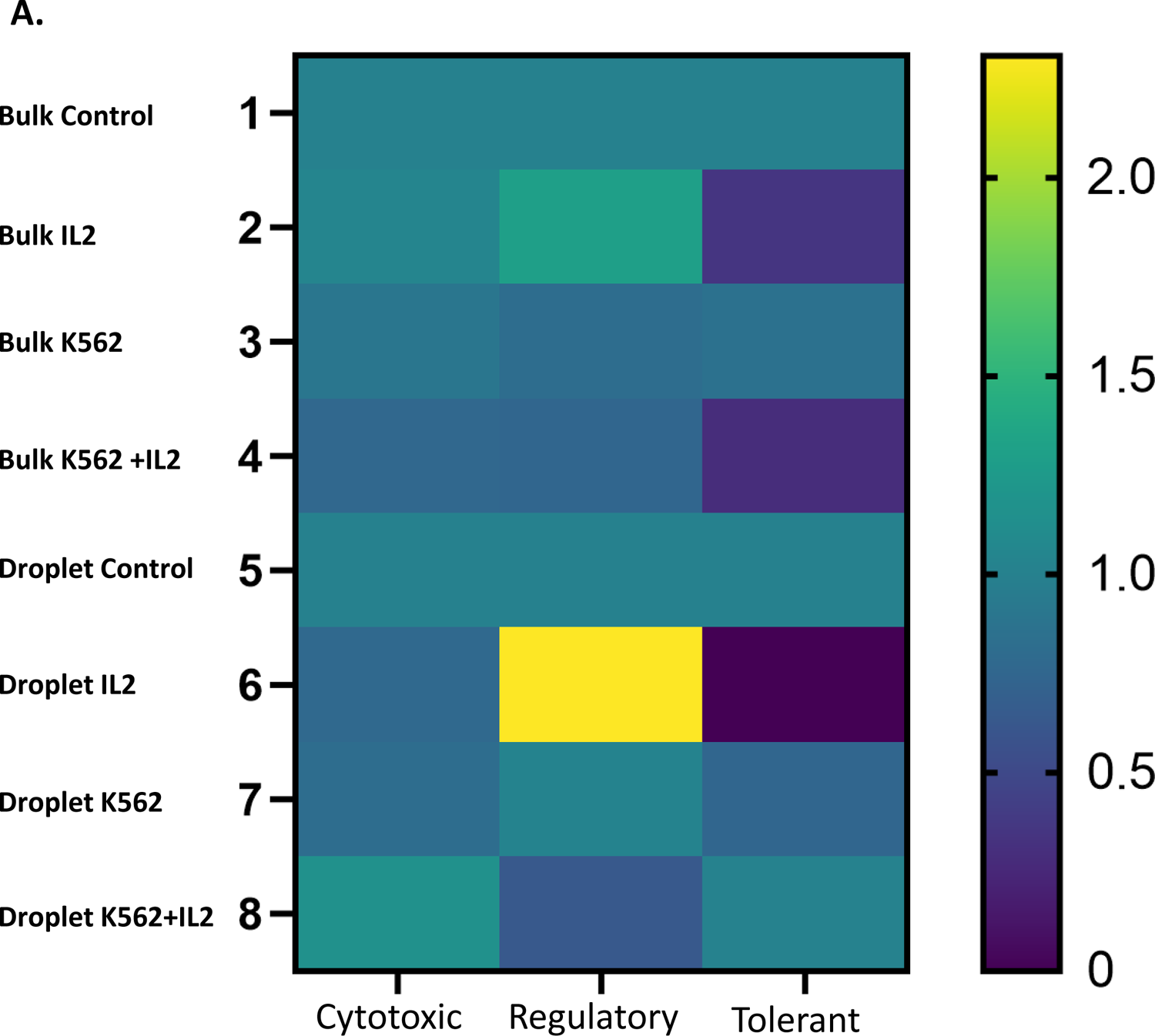
**A.** Heat map demonstrating the fold increment of different markers in NK cell with and without stimulation in bulk and droplets

**Supplementary Figure 3:**
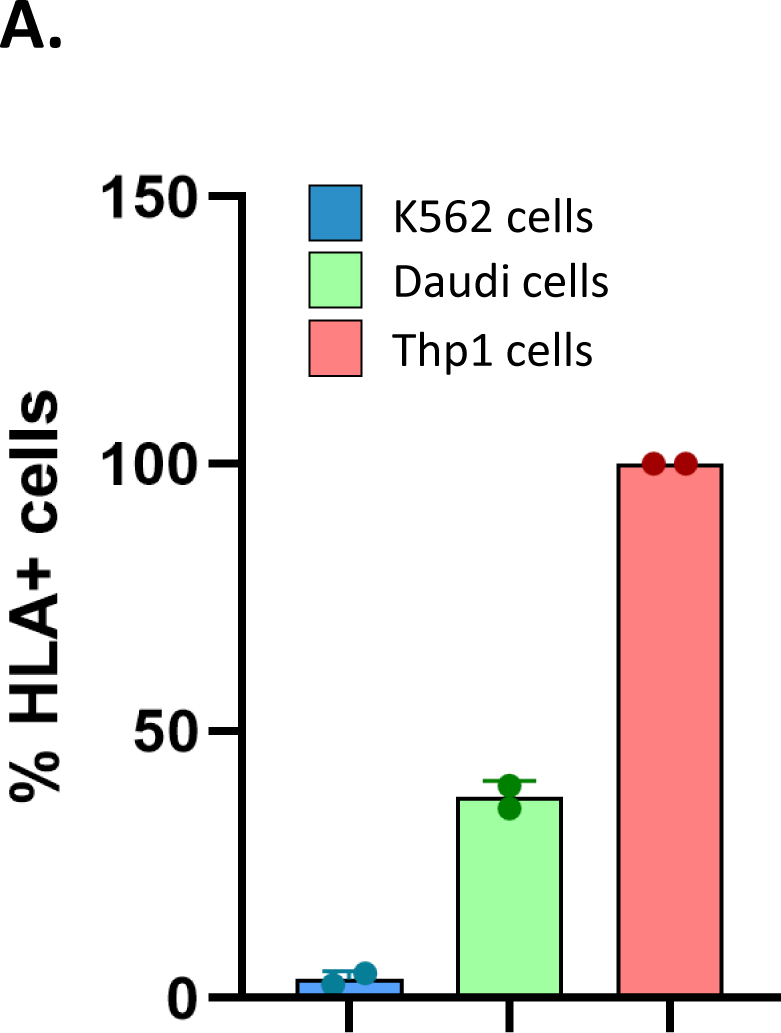
**A.** Expression of MHC-1 molecule by K562 (red), Daudi (green) and Thp1 cells (red); n=2 error bar represents standard error of mean.

**Supplementary Figure 4:**
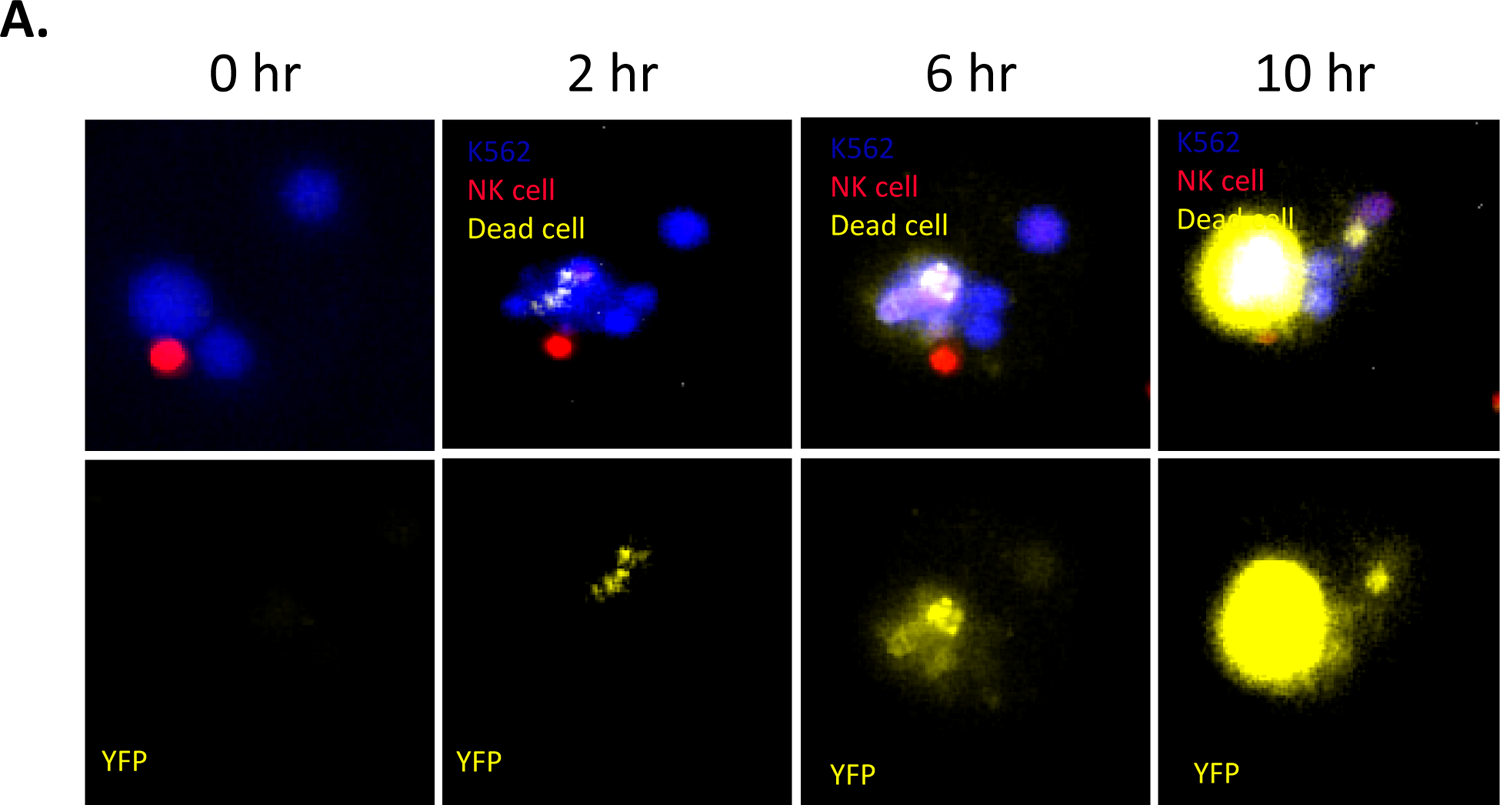
**A**. microscopic view of a single NK cell lysing 3 different K562 cells over 10 hours time period. Blue: K562 cell, Red: NK cell, Yellow: Dead cell

**Supplementary Figure 5:**
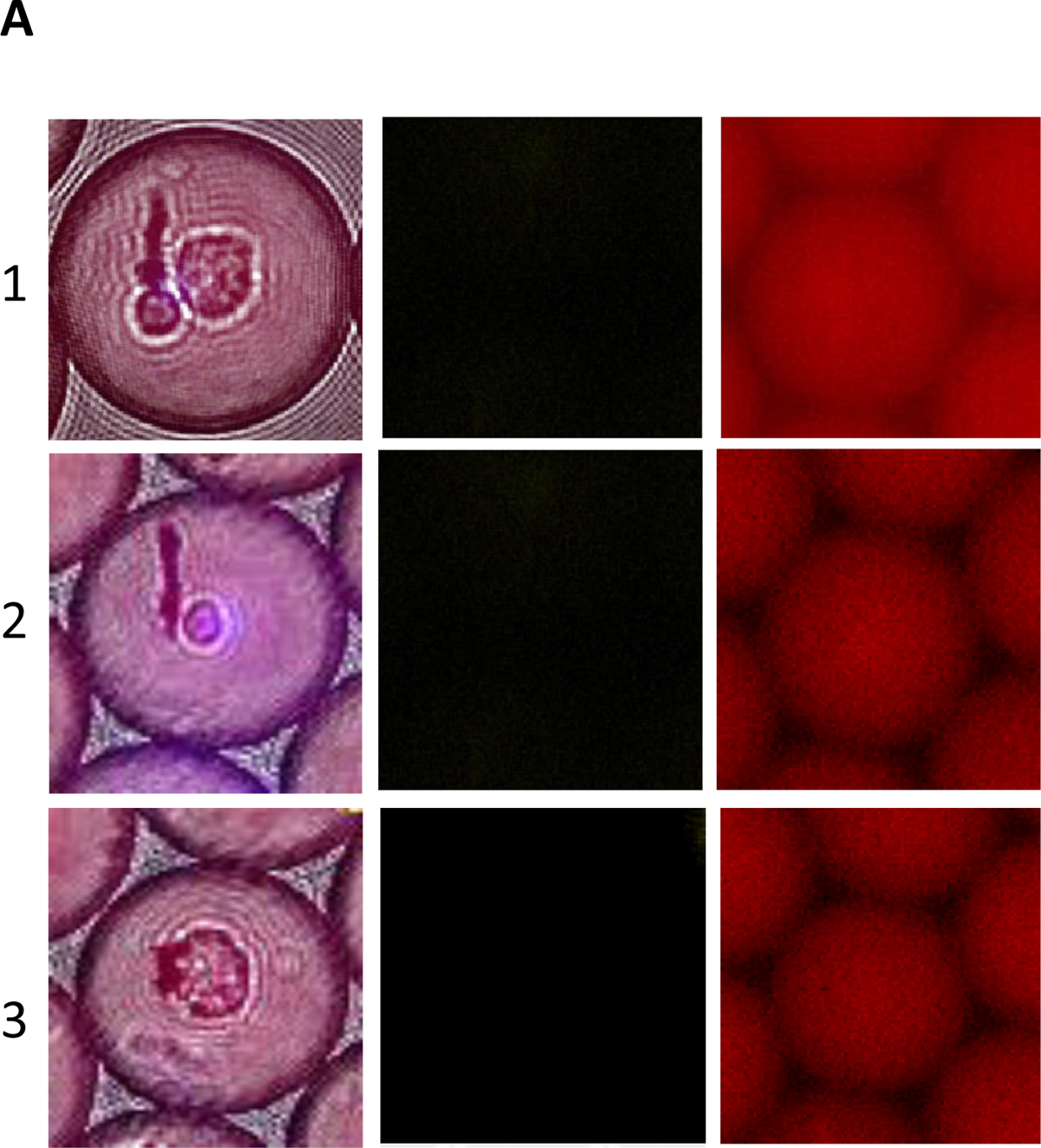
**A.** microscopic view showing different conditions in combinatorial assay for cytotoxicity and IFNγ secretion. 1. NK-K562 with no positive signal for both lysis and secretion. 2. only NK with no positive signal for secretion. 3. only K562 cells with no positive signal for cytotoxicity and secretion

